# Cell type-specific proteogenomic signal diffusion for integrating multi-omics data predicts novel schizophrenia risk genes

**DOI:** 10.1101/2020.05.28.121517

**Authors:** Abolfazl Doostparast Torshizi, Jubao Duan, Kai Wang

## Abstract

Accumulation of diverse types of omics data on schizophrenia (SCZ) requires a systems approach to jointly modeling the interplay between genome, transcriptome and proteome. Proteome dynamics, as the definitive cellular machinery in human body, has been lagging behind the research on genome/transcriptome in the context of SCZ, both at tissue and single-cell resolution. We introduce a Markov Affinity-based Proteogenomic Signal Diffusion (MAPSD) method to model intra-cellular protein trafficking paradigms and tissue-wise single-cell protein abundances. MAPSD integrates multi-omics data to amplify the signals at SCZ risk loci with small effect sizes, and reveal convergent disease-associated gene modules in the brain interactome as well as more than 130 tissue/cell-type combinations. We predicted a set of high-confidence SCZ risk genes, the majority of which are not directly connected to SCZ susceptibility risk genes. We characterized the subcellular localization of proteins encoded by candidate SCZ risk genes in various brain regions, and illustrated that most are enriched in neuronal and Purkinje cells in cerebral cortex. We demonstrated how the identified gene set may be involved in different developmental stages of the brain, how they alter SCZ-related biological pathways, and how they can be effectively leveraged for drug repurposing. MAPSD can be applied to other polygenic diseases, yet our case study on SCZ signifies how tissue-adjusted protein-protein interaction networks can assist in generating deeper insights into the orchestration of polygenic diseases.

## Introduction

The emergence of omics technologies has revolutionized neuropsychiatric research^1^ by generating high-throughput genomic data, bridging genome and transcriptome to phenome^2^. For example, genome-wide association studies (GWAS) such as Psychiatric Genomics Consortium (PGC)^3^ and CLOZUK consortium^4^ have created a repertoire of thousands of samples worldwide, leading to the discovery of many common variants associated with schizophrenia (SCZ). While such studies mark important milestones in schizophrenia research, they face critical challenges with regard to extracting novel biological insights and finding additional therapeutic targets or pathways. In fact, only one recognized drug target dopamine receptor D2 (*DRD2*) has been re-identified by GWAS^5^. It is not trivial to accurately pinpoint the corresponding risk genes in each GWAS risk locus, as such loci may cover a myriad of genes while the genuine causal variants may be far away from the top-ranking SNPs ^6^.

In addition to genetic association studies, tremendous efforts have been made over the course of years to understand the machinery of gene regulation. Whole-body proteomics data, such as the Human Protein Atlas^7,8^, now delineates protein expression not only across tens of various tissues but at certain cell-types, while drawing their subcellular localization. Moreover, large-scale epigenomics data such as Functional Annotation of the Mammalian Genome 5 (FANTOM5)^9^ and genome-scale chromosome conformation capture (Hi-C)^10,11^ technology have brought about unprecedented opportunities to elucidate long-range interactions among genetic loci. Given that individual omics data serve as complementary elements to each other, integrating multi-omics data-types can strengthen subtle disease signals from risk genes^5,12^. In fact, such multi-omics perspective amplifies signals from genetic loci with small effect sizes, and help support converging evidence on certain biological processes. This is of paramount importance in understanding polygenic diseases such as SCZ.

The current available omics data on SCZ are predominantly related to those of nucleic acids, e.g., genomics, transcriptomics, and epigenomics, while the use of proteomics information is quite limited ^13^. As the functional machinery in a cell, proteins essentially reflect the functional consequences of genome, epigenome, and transcriptome. Although proteins are treated as proxies of gene functions, multiple lines of evidences report a maximum of 60% correlation between the gene and protein expression levels in certain organisms^14,15^. Moreover, functionality of proteins is not restricted to their abundances, where other determinants such biochemical and physical properties such as subcellular localization, protein-protein interactions, and post-translational modifications affect such functions^16^. This mandates an inclusive in-depth analyses of the proteome and its physical and biochemical properties, not only at the tissue level but at the cell resolution in SCZ. Although proteomic investigations have been historically hampered due to the lack of low-cost and reliable high-throughput assay platforms^17,18^, there have been recent advances in improving the mass spectrometry-based proteomics platforms^19,20^ which has resulted in the generation of valuable resources such as the Human Protein Atlas^7,8^. On the other hand, subcellular fraction allows probing enrichment of proteins in micro-domains within cells (such as neurons), and offers insights into understanding the intracellular trafficking trajectories of proteins. There have been several proteomic studies on SCZ^21-24^ which mainly focus on observing the differential expression of proteins in post-mortem brains, without taking into account tissue- or cell-specific biochemical and biophysical interactions. For a full review on proteome studies in SCZ, refer to the reference^13^.

In this study, we introduce MAPSD, Markov Affinity-based Proteogenomic Signal Diffusion, a multi-omics network-based computational method to identify novel risk genes for polygenic diseases. MAPSD leverages multiple layers of omics information, as well as the under-studied proteome subcellular localization patterns and tissue-wise cell-specific abundances of proteins in tens of different tissues and a wide-range of cells, followed by propagating the biological signals across human interactome to characterize potential disease-associated risk genes. The proposed model has several unique advantages including: (a) it employs protein trafficking information in subcellular micro-domains in over 130 tissues and cell-types, including multiple regions in the brain from the Human Protein Atlas^7,8^, (b) MAPSD employs five layers of omics data including differentially expressed (DE) genes^2^, GWAS hits^3,4^, rare and *de novo* mutations^25^, differentially methylated genes^26-28^ and open chromatin accessibility data^29^; (c) MAPSD can effectively model interactions of genome, epigenome, transcriptome, and proteome at a single-cell resolution. Although we used SCZ as a test case in the study, MAPSD is flexible and can be effectively applied to other polygenic diseases other than SCZ. The outcome of MAPSD is accurate prediction of risk levels of all human genes in SCZ, which has led to the identification of a set of new candidate genes for SCZ. Our functional evaluation on these candidate genes indicate how the MAPSD-identified genes are predominantly enriched in certain cell-types within specific brain regions. In particular, the novel candidate genes identified by us are enriched for the targets of approved drugs for neurological disorders and suggest opportunities for re-purposing the existing therapies for SCZ.

## Results

### Overview of the MAPSD framework

MAPSD is a multi-step tissue/cell-specific proteogenomic method to identify risk genes through leveraging complementary biological signals from distinct omics data modalities. The overall structure of MAPSD is provided in **Figure 1**. MAPSD starts with a large-scale protein-protein interaction (PPI) network which was assembled from multiple sources^30-33^ (see Experimental Procedures). Using the PPI network, an affinity matrix is created. This matrix is binary in which if two nodes (proteins) interact then their corresponding matrix elements will be 1, otherwise 0. The PPI network is then adjusted to include molecular trafficking patterns. This adjustment is conducted using the subcellular localization data from the Human Protein Atlas (**Figure 2a**). The rationale behind this adjustment is that if two proteins being connected in the PPI network co-localize in the same micro-domain within the cell, then they are more likely to be interacting with each other. In total, 32 micro-domains have been used in this study. Therefore, the weight of connecting edges of co-localized proteins in the PPI network is amplified by a factor of 1.5 while the remaining edges have a weight of 1 (see Experimental Procedures). Using the adjusted affinity matrix, the Markov Transition Distribution Matrix *M* is created. Using Graph Laplacian concept in Graph Theory, a one-step probability distribution from each node to its neighbors is computed (see Experimental Procedures).

**Fig. 1.**
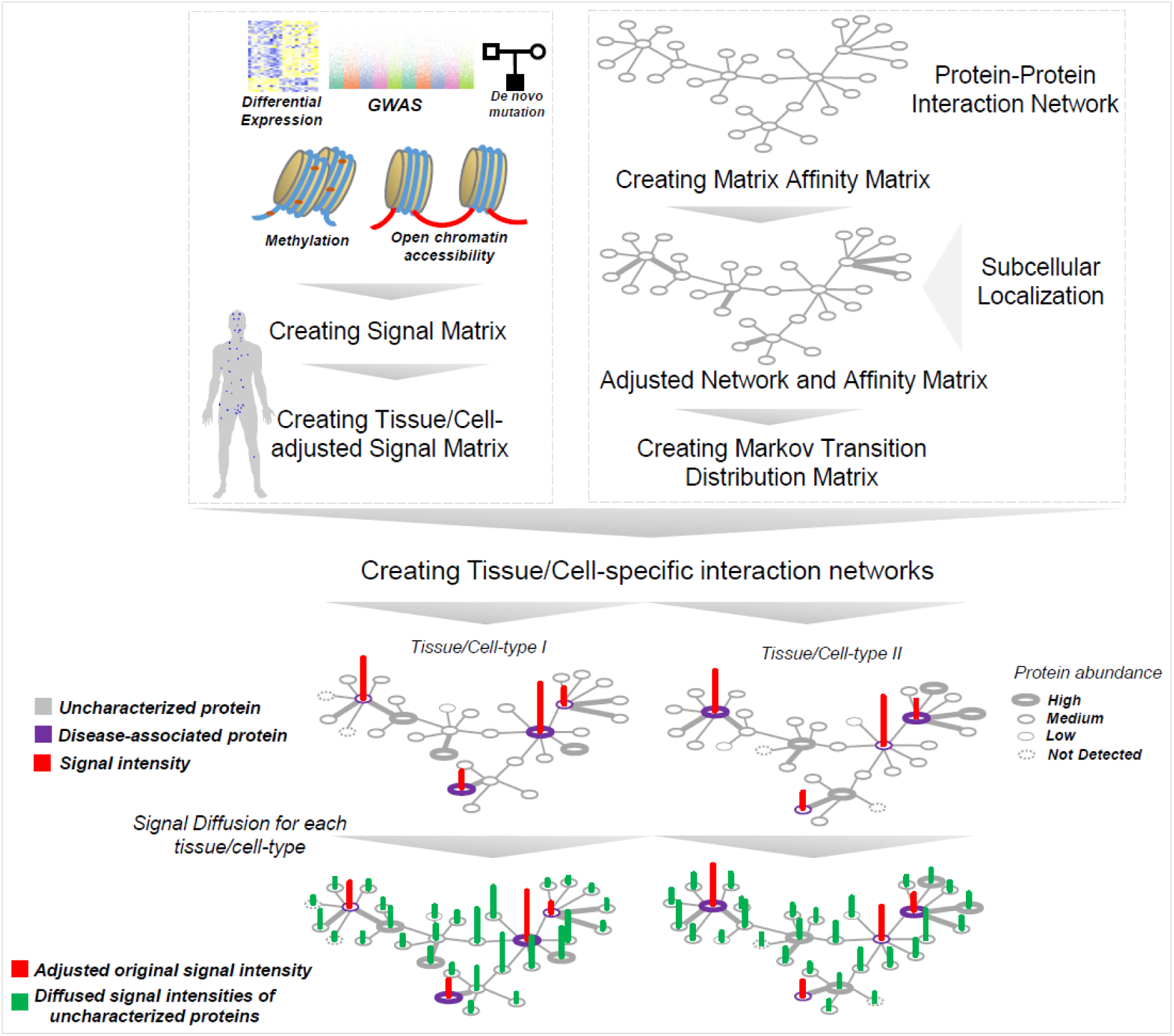
The structure of MAPSD. MAPSD steps include: creating the protein-protein interaction network followed by adjusting it for subcellular localizations, creating the Markov transition distribution matrix, assembling SCZ signatures from genome, epigenome, and transcriptome sources followed by creating the signal matrix and adjust it for different tissues and cell-types within them, creating tissue-cell-specific interaction networks, and signal diffusion across all of the dedicated networks to measure the disease signal intensities in unannotated proteins. Each dot on the human body scheme denoted the tissue being evaluated.

**Fig. 2.**
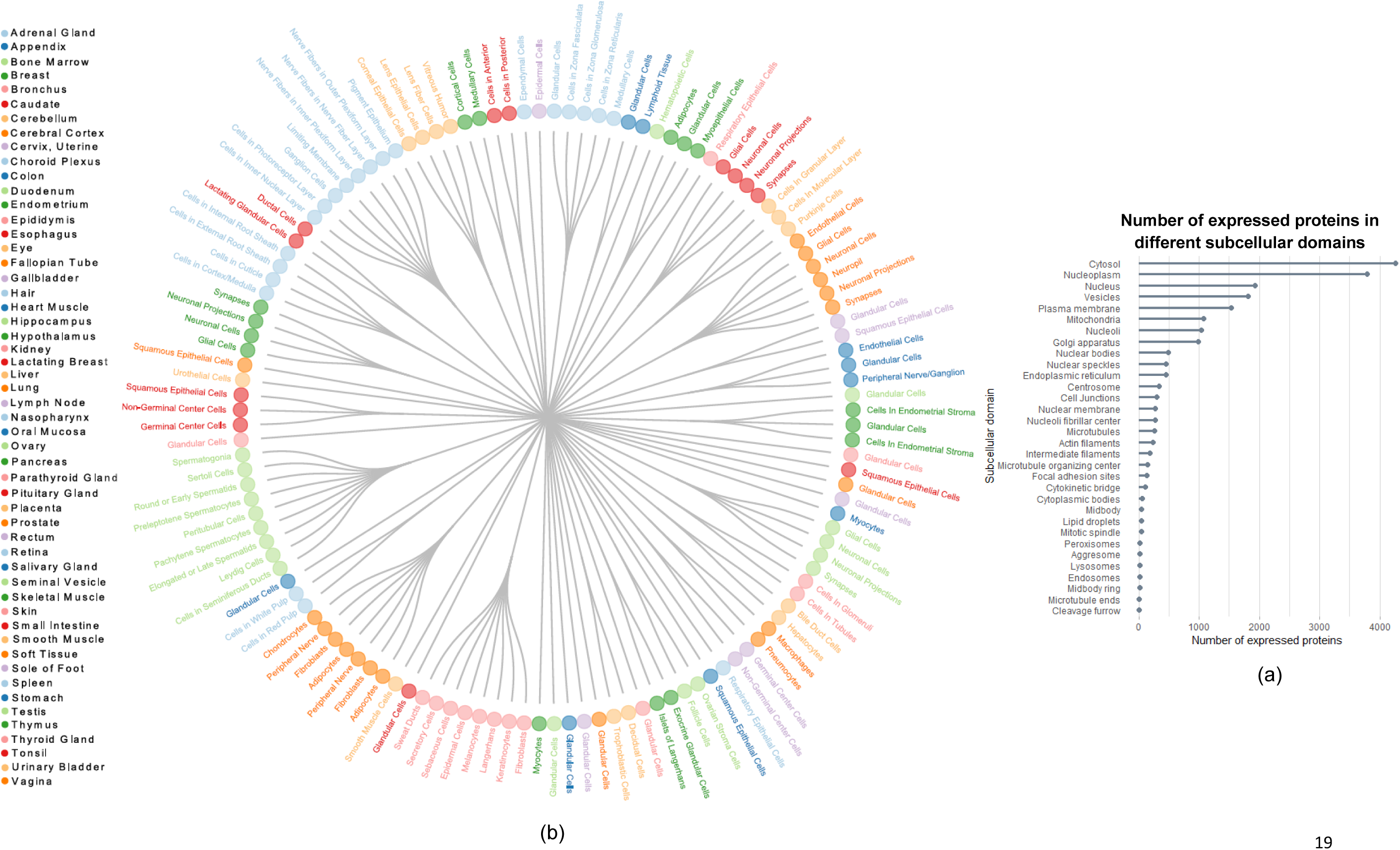
The list of cell-types and tissues used in this study. (a) 131 combination of cell-types and tissues. Each color denotes a tissue and the forks for each color represents their corresponding cell-types in this study; (b) The list of subcellular domains in this study followed by the number of proteins being expressed in each subcellular domain.

The multi-omics data sets have been collected from multiple sources (see Experimental Procedures). We used SCZ as a test case in our study to evaluate the MAPSD approach, due to the availability of large-scale genomics, transcriptomics and epigenomics data sets on SCZ. Five layers of omics data have been employed in this study including: differentially expressed (DE) genes, GWAS hits, rare and *de novo* mutation loci, differentially methylated loci, and loci being differentially accessible in open chromatin regions in neuronal cells. The corresponding Ensembl IDs for all of these loci were obtained and the final Signal Matrix was created. Since MAPSD operates at the single cell resolution, it needs to adjust the created initial signal vector *S* based on the tissues as well as their corresponding cell-types to project the variations between the protein abundances among them (see Experimental Procedures). To illustrate elements of the vector *S*, suppose a gene to be differentially expressed and differentially methylated in SCZ compared to controls. Then, the initial signal intensity of this gene in *S* equals 2. Using the available protein abundance data in various tissues and cell types from the Human Protein Atlas, we adjusted the signal vector *S* for 131 combinations of tissues and cell-types (**Figure 2b**, see Experimental Procedures). For instance, we have five regions in the brain including cerebral cortex, cerebellum, caudate, hippocampus, and hypothalamus as well as seven cell-types including neuronal cells, Purkinje cells, glial cells, endothelial cells, neutrophils, and cells in granular and molecular layers. Protein abundances vary across tissues and cell-types. Therefore, it is required to overlay the knowledge on such expression patterns onto the signal matrix *S*. The adjusted signal matrix is called *S*^*^ which shows the signal intensities of SCZ risk genes in all of the considered tissues and cell-types. In the next step, using the Markov operator matrix *M* and the created tissue/cell-specific signal intensity matrix *S*^*^, MAPSD diffuses the available adjusted signal intensities onto the adjusted networks aimed at estimating the disease signal intensities of the unknown proteins (see Experimental Procedures). Upon termination of the algorithm, MAPSD outputs the signal intensities of all of the proteins in 131 different combinations of tissues and cell-types, on which we conducted several tests. The MAPSD results are unbiased given that the adjusted network for signal diffusion is independently created from SCZ signal intensities and does not contain any prior information of the disease. Given that the PPI network is adjusted for subcellular localization of the nodes, the overall topology of the network shows a more realistic picture of subcellular molecular trafficking and protein interactions. The lower panel in **Figure 1** represents a toy example of diffused signals as well as the original SCZ signal intensities in two different cell-types. Given the abundance of proteins in each tissue and cell-type, the overall diffusion patterns of SCZ signals varies in the two networks.

### Applying MAPSD on SCZ to identify disease risk genes

We created a large PPI-network containing 232,801 edges and 16,185 nodes. As described above, considering five layers of omics evidences (gene expression, methylation, GWAS hits, rare and *de novo* mutation loci, and open chromatin regions), 3,915 genes were curated to be associated with SCZ with various degrees of signal intensities (**Figure 3a**). *DGKZ* showed the highest signal intensity of 4. *DGKZ* is a well-studied SCZ risk gene evidenced to be DE^2^ and differentially methylated^27^ as well as harboring GWAS hits^3,4^ and *de novo* mutations^25^. Six genes were found to have a signal intensity of 3 including *DNAJA4, TCF4, CHRNA2, CPNE8, GRIN2A*, and *ZNF536*. Notably, in a recent study^34^ we had identified *TCF4* to act as a transcriptional master regulator in SCZ, based on expression network analysis of human dorsolateral prefrontal cortex (DLPFC). Upon initiating the diffusion process, MAPSD terminated the diffusion at the time step *t* = 3 (**Figure 3c**). A sharp decrease in **Figure 3c** indicates the tendency of the graph toward over-smoothness. Therefore, *t* = 3 is an appropriate cut-off point to prevent this phenomenon. After completion of the diffusion process, we sought to check how many of the SCZ risk genes show the highest signal intensity in all of the brain regions (**Figure 3b**). We can see that *DGKZ* as well as two other genes with signal intensity of 3 were preserved in the brain. MAPSD resulted in 704 genes (4.4% of the total) to bear the highest SCZ risk signal uniquely in several brain regions, including cerebral cortex, cerebellum, hippocampus, and caudate. We checked this gene set to look for the SCZ risk genes (which were used as the input to the method) showing the highest risk signal intensity upon executing the MAPSD. We found that 190 genes have the highest signal intensities only in the brain (**Figure 3b**). We checked the signal intensity of the remaining SCZ-associated genes (n=3,725). We found 3,480 genes to bear the highest signal intensity in the brain as well as at least one other tissue other than the brain, while 245 genes showed higher risk signals in other tissues other than brain.

**Fig. 3.**
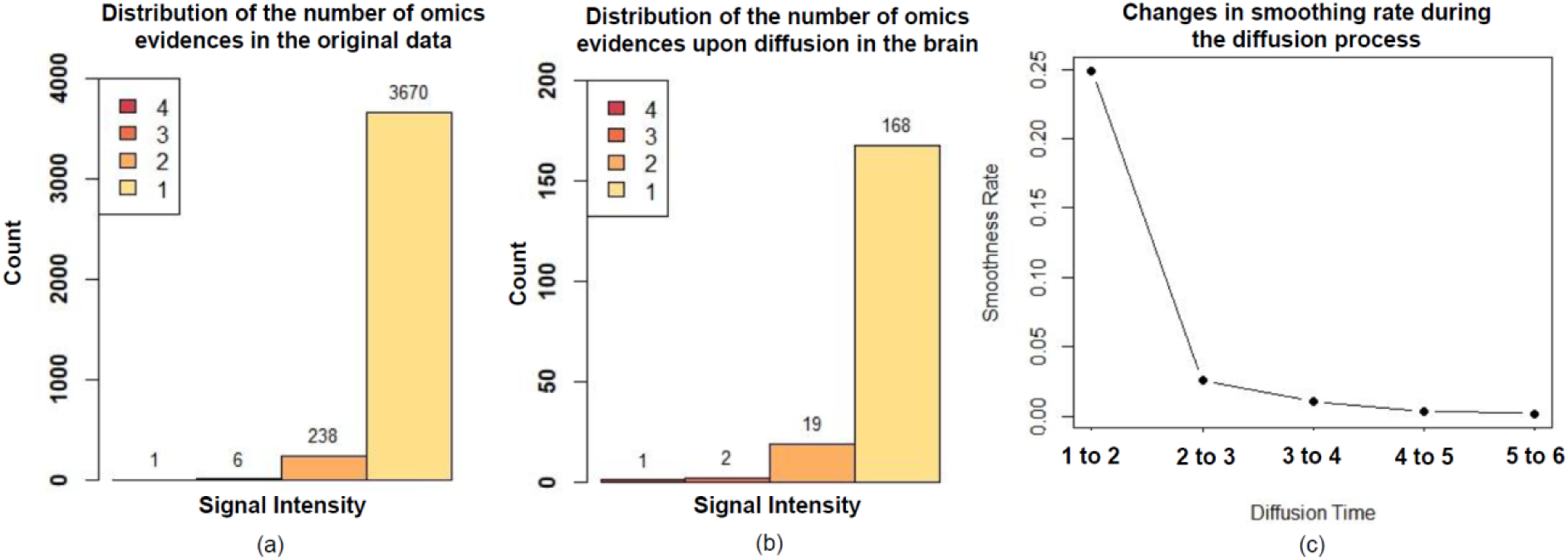
Distribution of SCZ signal intensities. (a) Distribution of initial signal intensities in the original signal vector; (b) Distribution of initial signal intensities enriched in the brain after signal diffusion; (c) Changes of smoothing rate during the diffusion time.

### MAPSD-identified SCZ risk genes are enriched in specific subcellular domains in neuronal cells

To evaluate the reliability of the MAPSD-identified candidate risk genes, we separated the 704 identified genes with the highest signal intensity in the brain into two groups: 190 known SCZ-risk genes and 514 newly identified genes (**Figure 4a-b**). Using the protein abundances from the Human Protein Atlas, we checked in what specific brain regions and cell-types the protein products of these genes are expressed. Of 190 known SCZ risk genes, 126 genes (66.3%) were highly expressed in neuronal cells in cerebral cortex while in total, 138 genes (∼72.3%) of the entire gene set were highly expressed in various cells-types in cerebral cortex. We next sought to evaluate the set of newly identified genes in the brain. We made a similar analysis on the 514 newly identified gene set by MAPSD. Among them, 360 genes (∼70%) were highly expressed in neuronal cells in cerebral cortex. In total 396 genes were highly expressed only in cerebral cortex which accounts for 77% of the total number of the newly identified gene set. Notably, these observations reveal an agreement between the enrichment patterns of the both gene sets and suggests reliable cell-specificity of the MAPSD approach. This finding is in agreement with the cell-types suggested to be underlying SCZ pathogenesis^35^. In an important study, Skene *et al.*, had investigated the enrichment of SCZ common variants in adult brain temporal cortex and prefrontal cortex. Cell-types being studied in these regions included: astrocytes, oligodendrocyte progenitor cells (OPCs), oligodendrocytes, microglia, pyramidal neurons, and cortical interneurons. In both regions, pyramidal neurons and interneurons shared the highest degree of enrichment of GWAS loci compared to the other cell-types. Our observations also show that the identified risk genes, at protein level, are predominantly highly expressed in neuronal cells compared to other available cell-types in this region. We also noted that endothelial cells share the lowest fraction of SCZ risk genes in our study. This is also the case in the findings of Skene *et al.*, in which the enrichment of SCZ common variants in endothelial cells in prefrontal cortex is the lowest compared to the other cell-types.

**Fig. 4.**
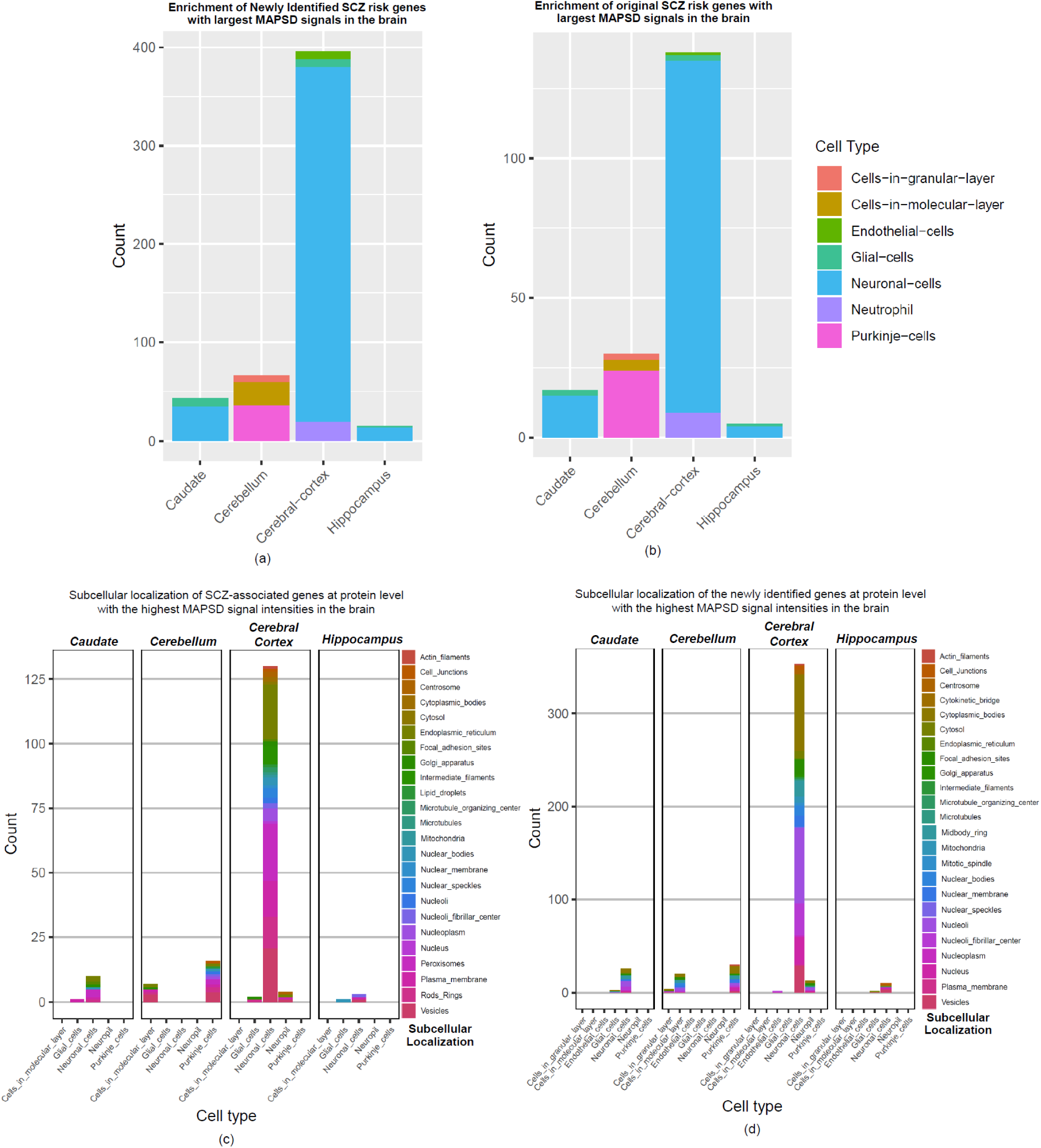
Expression patterns of MAPSD brain-specific genes at cell resolution and subcellular domains. (a) Frequency of MAPSD newly identified SCZ risk genes at single cell resolution to be highly expressed in four brain regions; (b) Frequency of MAPSD original SCZ risk genes at single cell resolution to be highly expressed in four brain regions; (c) Frequency of MAPSD original SCZ risk genes at protein level to be highly expressed in various subcellular domains in five cell-types across four different brain regions; (d) Frequency of MAPSD newly identified SCZ risk genes at protein level to be highly expressed in various subcellular domains in five cell-types across four different brain regions.

We were interested in finding the localization of SCZ risk genes in subcellular domains, using the subcellular localization domains obtained from Human Protein Atlas (**Figure 2b**). An immediate observation is significant enrichment of SCZ risk loci at protein level in various sub-cellular micro-domains of neuronal cells within cerebral cortex (**Figure 4c**). 78% of the original SCZ risk genes found by MAPSD were enriched in neuronal cells in cerebral cortex and across different subcellular domains. Among them, ∼96% were enriched only in neuronal cells across different micro-domains. Further focusing on neuronal cells, we found that five micro-domains including cytosol, nucleus, nucleoplasm, plasma membrane, and vesicles share ∼70% of the entire SCZ-associated protein products in cerebral cortex. Across the entire subcellular micro-domains, cerebellum harbors ∼13% of the candidate SCZ-risk genes, in which Purkinje cells shares the highest fraction of SCZ candidate risk genes at protein level.

We compared the enrichment patterns of the MAPSD newly identified genes with the known SCZ risk genes based on their corresponding micro-domains. Similar to the SCZ risk genes, subcellular micro-domains in neuronal cells within cerebral cortex share the largest fraction of the identified genes. We checked the newly identified gene set in cerebral cortex. Considering all of the micro-domains, ∼96% of the entire identified proteins are expressed predominantly in neuronal cells (**Figure 4d**). Within neuronal cells, five micro-domains share 72.5% of these proteins including: cytosol, nucleus, nucleoplasm, plasma membrane, and vesicles. This fraction is very similar to the localization of SCZ-associated protein products in neuronal cells within cerebral cortex.

We compared the proportions of enrichment of SCZ genes and the identified genes based on their localizations within each cell in separate brain regions. In cerebral cortex, considering all of the micro-domains and cell types, fractions of the both known SCZ risk genes and MAPSD newly identified genes were similar with no significant difference observed (Chi-Square P-value=0.79). We further compared the differences between the proportions of the major subcellular domains indicated above in neuronal cells within cerebral cortex. Except Vesicles (Chi-Square P-value=0.018), no significant difference was observed between their proportions: plasma membrane (Chi-Square P-value=0.9432), cytosol (Chi-Square P-value=0.114), nucleus (Chi-Square P-value=0.842), nucleoplasm (Chi-Square P-value=0.191). These observations extends further support regarding efficacy of MAPSD in modeling a more realistic map of proteomic properties of SCZ at the cellular resolution.

### MAPSD recovers potential disease-associated susceptibility protein complexes

In addition to finding novel candidate risk genes, MAPSD can also reveal protein complexes that may be involved in disease pathogenesis. We tested MAPSD to show how it can facilitate recovering the SCZ risk signals in the brain. We ran MAPSD 100 times and in each time randomly removed one SCZ risk gene with the highest signal intensity in the brain. MAPSD successfully recovered their signal intensities to bear the highest SCZ signal intensities in the brain. As an example, we illustrate the signal intensity of two SCZ risk genes (*DGKZ* and *ST8SIA2*) to show the highest signal intensity levels in the brain. *DGKZ* has been implicated in SCZ to be DE, differentially methylated, as well as harboring *de novo* mutations. MAPSD signal intensities for this gene (**Figure 5a**) are the highest in three regions including: neuronal cells in cerebral cortex, Purkinje cells in cerebellum, and neuronal cells in caudate. *ST8SIA2* (**Figure 5b**) is known to be associated with SCZ in various ways such as its impacts on cerebral white matter diffusion properties in SCZ^36^ as well as harboring multiple SCZ-associated single nucleotide polymorphisms (SNPs)^3,37^. After removing this gene from the initial signal matrix, we ran MAPSD and observed that MAPSD yields the highest SCZ signal intensities in cerebral cortex and cerebellum. These experiments verifies the robustness of MAPSD when the initial signal information for a disease is partially complete and that the method is capable to re-identify genuine SCZ risk loci given the topology of the adjusted PPI networks as well as proteome information incorporated into the model. Looking at the outcome of MAPSD in the newly identified gene set, we found several genes to be implicated in other neurological disorders. Considering that MAPSD can recover known SCZ-associated risk factors, we hypothesize that the newly identified genes may potentially implicate in SCZ. On the other hand, we are already aware that many psychiatric disorders such as SCZ, autism, and bipolar disorder share substantial genetic susceptibility ^38^. Therefore, as a proof of concept, we picked some of the top MAPSD genes with the highest signal intensity and evaluated whether they have already been indicated in other brain diseases.

**Fig. 5.**
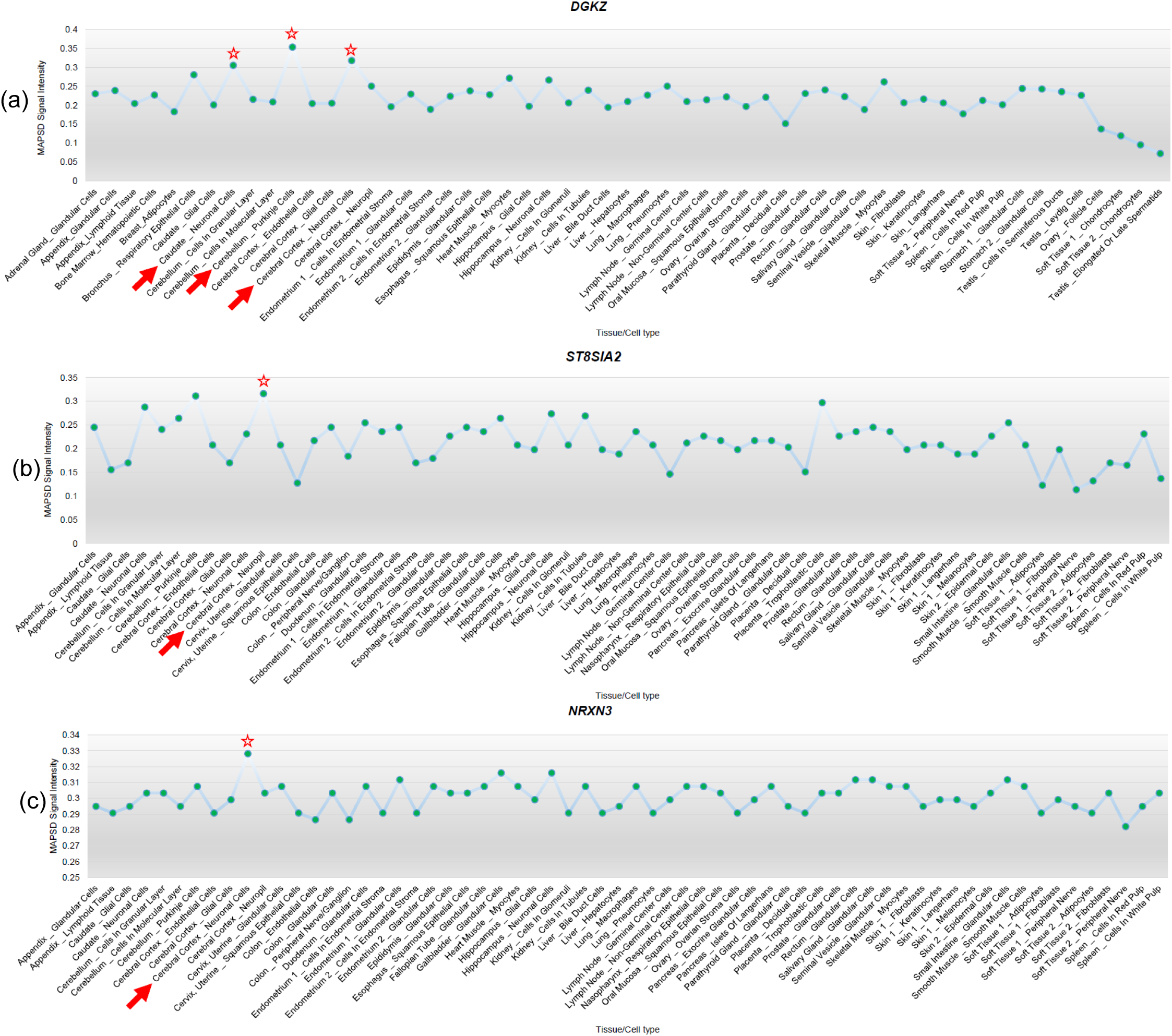
MAPSD signal intensities upon diffusion in three genes. (a) MAPSD signal intensities of the SCZ risk gene DGKZ; (b) MAPSD signal intensities of the SCZ risk gene ST8SIA2; (c) MAPSD signal intensities of the gene DGKZ NRXN3 found to show the highest risk signals in the brain.

As a proof of concept, we picked *NRXN3* which shows the highest signal intensity in neuronal cells in cerebral cortex upon executing MAPSD (**Figure 5c**). The autism risk gene *NRXN3*^39,40^ is a member of Neuroxin gene family which encodes neuronal adhesion proteins with critical roles in synapse development and function. Although restricted evidences such as copy number variation^41^ and a polymorphism^42^ on *NRXN3* have been reported to be associated with SCZ in small population cohorts, its association to the disease has not been replicated ^43^ or widely recognized. We investigated the PPI network to look for the genes connected to *NRXN3. NRXN3* is directly connected to six genes, where the majority of them are significantly associated with diseases related to the central nervous system (CNS). These genes include: *NLGN1, NLGN2, NLGN3, CASK, AFDN*, and *PAX4. NLGN1, NLGN2*, and *NLGN3* belong to the family of neuronal cell surface proteins, Neuroligin, and are involved in formation of CNS synapses^44^. They have been implicated in epilepsy^45^, autism spectrum disorders (ASD)^46^, and post-traumatic stress disorder (PTSD)^47^. Notably, MAPSD recapitulated these three genes in the brain where *NLGN1* and *NLGN2* were input to the model as SCZ risk genes yet *NLGN3* was identified by MAPSD as a susceptibility disease risk gene. This finding is in concordance with the well-established observations that Neuroligin protein members act as ligands for Neuroxins, resulting in the connections between neurons and generation of synapses^48^. *CASK* and *AFDN* have also been implicated in CNS diseases such intellectual disabilities^49,50^ and CNS leukemia^51,52^, respectively. Given that *AFDN* interacts with *NRXN3*^*53*^, we can conclude that MAPSD is capable of recovering high-risk loci in protein complexes and can infer a big picture of the converging disease-risk modules in the human interactome.

### Tissue and developmental stage-specific expression of MAPSD risk genes

To further gain evidence supporting their disease relevance, we analyzed the tissue-specific expression levels of the identified SCZ risk genes at mRNA level. For this, we used gene expression levels on 53 different tissues from the Genotype-Tissue Expression (GTEx) project^54^. GTEx data contains mRNA levels across the entire transcriptome, which enables specifying to what extent a gene is expressed in distinct tissues. We divided the MAPSD risk genes into two groups, including the known SCZ risk genes with the highest signal intensities in the brain and newly identified genes with the highest signal intensity in the brain. We queried the GTEx data and observed that in both sets, the outputs of MAPSD are highly enriched in brain tissues (**Figure 6a**). In fact, frontal cortex showed remarkably higher enrichment scores, which is supported by the previous findings regarding its implications in SCZ^2,55^. The extent of enrichment in distinct brain regions were different. For instance, frontal cortex and cerebral hemisphere represented a much stronger enrichment significance compared to other regions in the brain, while amygdala and hippocampus, despite being significant, were less implicated in our analysis. In addition to the provided significance P-values, we calculated the fold enrichment ratios (FER) for the top five significant brain regions for the set of identified genes including: frontal cortex (FER=8.9), cortex (FER=8.8), anterior cingulate cortex (FER=21.7), nucleus accumbens (FER=5.1), and cerebellar hemisphere (FER=2.9). These observations suggest that integrating cell-specific genome and proteome knowledge in modeling the disease can lead to more sensitive and reliable identification of novel risk factors.

**Fig. 6.**
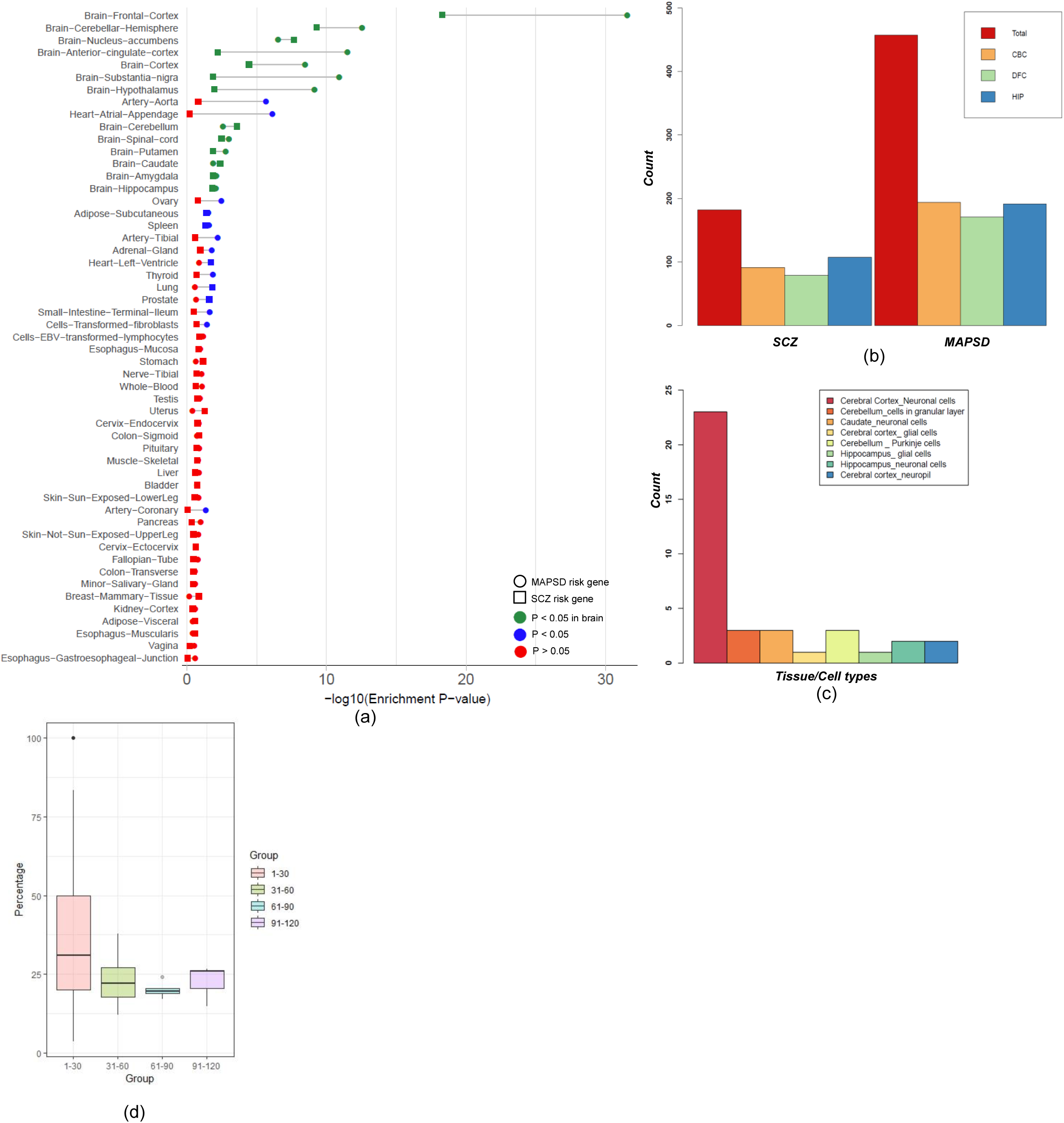
Tissue-wise enrichment statistics for SCZ and MAPSD identified genes at gene expression level. (a) –log10(P-value) of SCZ and MAPSD risk genes with the highest signal intensity in brain tissues in GTEx consortium gene expression data; (b) The number of differentially expressed SCZ and MAPSD risk genes in cerebral cortex (CBC), dorsolateral frontal cortex (DFC) and hippocampus ^57^ between prenatal and postnatal developmental stages using BrainSpan data; (c) Number of MAPSD risk genes to be the targets of FDA-approved drugs being enriched in specific cell-types in certain brain regions; (d) Percentage of SCZ-associated genes to be direct neighbors of the MAPSD identified genes where each color represents MAPSD genes with a certain number of immediate connecting nodes in the PPI network.

Since SCZ is likely a neurodevelopmental disorder, we next investigated if the brain-specific MAPSD genes are dysregulated during various developmental stages in human brain. We used the Atlas of the Developing Human Brain (Brain Span)^56^ on three brain regions including dorsolateral frontal cortex (DFC), cerebral cortex (CBC), and hippocampus ^5757^. Next, we divided the data into two large categories of prenatal and postnatal stages, each with various time points. Prenatal stage includes: 0-12 post-conception weeks (pcw), 13-24 pcw, and 25-36 pcw. Postnatal stages include: 0-2 yr, 3-8 yr, 9-16 yr, and >17 yr. We averaged the expression levels of each MAPSD genes across different stages of pre- and postnatal stages and looked for DE genes (**Figure 6b**). Our observation indicates that almost half these genes were DE in postnatal stages versus the prenatal stages. The overall pattern of the number of DE genes in SCZ and MAPSD genes was almost similar. We were interested to specify what biological pathways are disrupted by the dysregulated genes during neurodevelopment in DFC, CBC, and HIP. We conducted pathway enrichment analysis (see Experimental Procedures) on these three gene sets. Although several pathways were nominally significant, none of them passed the false discovery rate (FDR) threshold of 0.05. On the other hand, checking the SCZ-associated genes which demonstrated the highest signal intensity while being DE during neurodevelopment led to finding multiple pathways that are statistically significant (FDR<0.05). The majority of these pathways were shared by the three regions such as: glutamatergic synapse (DFC: FDR=2.3×10^−8^, Fold Enrichment Ratio (FER)=15.4; CBC: FDR=8.5×10^− 9^, FER=13.8; HIP: FDR=2.8×10^−7^, FER=11.8), calcium signaling pathway (DFC: FDR=1.32×10^−7^, FER=10.4; CBC: FDR=8.5×10^−9^, FER=10; HIP: FDR=2.8×10^−7^, FER=8.7), circadian entertainment (DFC: FDR=7.5×10^−6^, FER=13.2; CBC: FDR=2.2×10^−7^, FER=13.6; HIP: FDR=4.4×10^−7^, FER=12.8), and cholinergic synapse (DFC: FDR=2.1×10^−5^, FER=11.3; CBC: FDR=7.3×10^−4^, FER=8.2; HIP: FDR=2×10^−4^, FER=8.8).

### MAPSD risk genes are potential drug targets

We were interested in whether the MPASD-identified SCZ risk genes act as targets of known drugs related to CNS. We used the list of FDA-approved drug targets by Santos *et al*.^58^ comprising 4,631 drug-target connections as well as their mechanism of action. The data contained 881 unique protein targets in which the Ensemble IDs of 713 proteins were obtained. Among 514 newly identified MAPSD risk genes, we found 38 genes (**Table 1**) to be the targets of available FDA-approved drugs (FET P-value=2.68×10^−4^). We found multiple calcium channel mRNAs to be of high risk signal intensities such as *CACNB1, CACNG2, CACNG3*, and *CACNG7*. These genes are known to be the targets of fragile X mental retardation protein (FMRP) which cause Fragile X syndrome (FXS) and autistic symptoms^59^. These proteins were highly enriched in the brain, specifically in neuronal cells in cerebral cortex (**Figure 6c**). We were interested in finding the genes which are already targets of drugs developed for CNS diseases. 21 (56%) of the 38 genes were targets of drugs developed for CNS-related diseases (**Supplementary Figure 1**). Some of these genes are well-documented risk loci in neurological diseases. For instance, *SCN1A*, a voltage-dependent sodium channel gene is known to be associated with epilepsy^60,61^. These genes are essential in generating action potentials in neurons and muscles. We found this gene to be the target of 16 drugs primarily developed to treat epilepsy. We had found this gene to exhibit the highest signal intensity in neuronal cells in cerebral cortex. Similarly, *SCN3A*, an epilepsy gene was picked up by MAPSD in neuronal cells in cerebral cortex and hippocampus. These two genes have been widely studied in epilepsy as well as mental retardation and other neuropsychiatric disorders^60^. We recognize that these genes may have a different mode of action (gain-of-function versus loss-of-function) in different brain disorders, but our analysis demonstrated a proof of principle that MAPSD may facilitate drug repurposing efforts by integrating more fine-grained (tissue-specific, cell type-specific, and subcellular localization-specific) omics information on brain disorders.

**Table 1.** MAPSD identified risk genes which are targets of available drugs.

| Gene | UniProt Accession | PROTEIN NAME |
| --- | --- | --- |
| CACNB1 | Q02641 | Voltage-dependent L-type calcium channel subunit beta-1 |
| CACNG7 | P62955 | Voltage-dependent calcium channel gamma-7 subunit |
| GRIA4 | P48058 | Glutamate receptor 4 |
| GRIA1 | P42261 | Glutamate receptor 1 |
| ADRA1B | P35368 | Alpha-1B adrenergic receptor |
| KCNA4 | P22459 | Potassium voltage-gated channel subfamily A member 4 |
| CA12 | O43570 | Carbonic anhydrase 12 |
| KCNQ2 | O43526 | Potassium voltage-gated channel subfamily KQT member 2 |
| NDUFB3 | O43676 | NADH dehydrogenase [ubiquinone] 1 beta subcomplex subunit 3 |
| NDUFS7 | O75251 | NADH dehydrogenase [ubiquinone] iron-sulfur protein 7, mitochondrial |
| GRIN2C | Q14957 | Glutamate receptor ionotropic, NMDA 2C |
| HCN4 | Q9Y3Q4 | Potassium/sodium hyperpolarization-activated cyclic nucleotide-gated channel 4 |
| IL12B | P29460 | Interleukin-12 subunit beta |
| CHRM1 | P11229 | Muscarinic acetylcholine receptor M1 |
| LAMC3 | Q9Y6N6 | Laminin subunit gamma-3 |
| CHRNA4 | P43681 | Neuronal acetylcholine receptor subunit alpha-4 |
| CACNG2 | Q9Y698 | Voltage-dependent calcium channel gamma-2 subunit |
| EPHB4 | P54760 | Ephrin type-B receptor 4 |
| PDE8A | O60658 | High affinity cAMP-specific and IBMX-insensitive 3',5'-cyclic phosphodiesterase 8A |
| SCN1A | P35498 | Sodium channel protein type 1 subunit alpha |
| CHRM2 | P08172 | Muscarinic acetylcholine receptor M2 |
| HRH1 | P35367 | Histamine H1 receptor |
| CNR1 | P21554 | Cannabinoid receptor 1 |
| GRIK1 | P39086 | Glutamate receptor ionotropic, kainate 1 |
| MT-ND2 | P03891 | NADH-ubiquinone oxidoreductase chain 2 |
| KCNQ3 | O43525 | Potassium voltage-gated channel subfamily KQT member 3 |
| GRIN3B | O60391 | Glutamate receptor ionotropic, NMDA 3B |
| PAH | P00439 | Phenylalanine-4-hydroxylase |
| GRIK3 | Q13003 | Glutamate receptor ionotropic, kainate 3 |
| HCRTR1 | O43613 | Orexin receptor type 1 |
| CACNG3 | O60359 | Voltage-dependent calcium channel gamma-3 subunit |
| KCNQ4 | P56696 | Potassium voltage-gated channel subfamily KQT member 4 |
| PTGER1 | P34995 | Prostaglandin E2 receptor EP1 subtype |
| TYMS | P04818 | Thymidylate synthase |
| SLC12A1 | Q13621 | Solute carrier family 12 member 1 |
| GNRHR | P30968 | Gonadotropin-releasing hormone receptor |
| HCN3 | Q9P1Z3 | Potassium/sodium hyperpolarization-activated cyclic nucleotide-gated channel 3 |
| SCN3A | Q9NY46 | Sodium channel protein type 3 subunit alpha |

Another highly connected gene within the created drug-target network was *HRH1*. This gene was found to be the target of 51 drugs, of which 10 were developed for CNS diseases. This gene showed the highest MAPSD signal intensity in neuronal cells in cerebral cortex despite not being used initially as a SCZ signature in MAPSD. A few studies have investigated its association with SCZ. For example, Nakai *et al*^*62*^ have shown the possible associations between *HRH1* and SCZ, despite of a borderline evidence for association in GWAS^63^. We found this gene to be connected to *ADRA1B* through two antipsychotic drugs Chlorpromazine and Trimipramine. Such interdependencies between the original SCZ risk genes supplied to MAPSD and the identified high signal genes further supports an orchestrated mechanism of the disease through interactions in convergent modules in the human interactome.

Among the identified genes to be drug targets, *CHRM1* and *CHRM2* were found to be targeted by over 30 drugs, 8 related to CNS. These genes are implicated in alcohol dependence^64^, major depression^65^, as well as possible involvements in SCZ^66^. In addition to the identified genes which might have been implicated in neuropsychiatric disorders, MAPSD revealed new candidates for treatment of SCZ. For instance *SLC12A1*, a solute carrier transporter, was found with the highest signal intensity in the brain to be targeted by five drugs. This gene is essentially targeted to reduce edema caused by kidney or heart failure. However, granted the role of such membrane-bound proteins in transferring substrates within the cell such as dopamine and serotonin^67^, they can be further studied for the treatment of SCZ.

## Discussion

In our view, the extreme polygenic nature of complex psychiatric disorders, such as SCZ, necessitates taking a more holistic view on the overall system of the diseases. One critical component of such a system is proteome and its dynamics, given that proteins are in fact work horses of intra-cellular activities. Proteins reflect the genetic, epigenetic, and transcriptomic alterations which are caused by the disease. Yet, research on proteome lags behind other omics data-types, especially those generated on DNA and RNA levels^13^, due to technical limitations in data generation. Recent advances in proteome experimental paradigms has created new horizons to further employ proteome knowledge in studying SCZ. Integrated analysis of omics data-types at nucleic acid and amino acid levels makes it possible to accurately pinpoint SCZ drivers as well as accurate isolation of gene modules whose orchestrated interactions may confer susceptibility to the disease. Taking a multi-layer approach to SCZ, we introduced MAPSD, a proteogenomic signal diffusion method which accounts for subcellular localization of the proteins and intra-cellular trafficking in an integrated manner. Our study demonstrated the effectiveness of the MAPSD in recovering known SCZ risk genes and identifying novel candidate risk genes, and in identifying possible drug targets for drug repurposing studies.

MAPSD has unique characteristics which are worth further discussion. MAPSD features modeling the protein localization in subcellular micro-domains as well as tissue-wise cell-specific distribution of protein abundances in the human body. Taking all this information into account, MAPSD creates dedicated cell-specific PPI network for tens of distinct human tissues. This allowed us to create more realistic PPI networks that can lead to more accurate prediction of disease drivers. MAPSD jointly uses GWAS hits, DE genes, rare and *de novo* mutations, and open chromatic accessibility data followed by diffusing this repertoire of information into each dedicated cell-specific PPI network to predict the signal intensities of novel candidate genes and their potential role in the disease onset and progression. The Markov Affinity-based criterion borrowed from Graph Theory as well as the designed termination criterion ensures accurate transition of information across the network, while avoiding over-smoothing the signal intensities. Therefore, the highest amount of information will flow through the network while preventing the signals at each node are distinctive enough. MAPSD enables ranking the genes related to SCZ given their signal intensity levels in the brain.

An important strength of MAPSD is that the identified novel disease risk gene may not by immediate neighbors of known SCZ risk genes. For example, 217 genes out of 514 (∼42%) identified risk genes by MAPSD are not directly connected to disease susceptibility loci. We checked the topology of the PPI network on the identified MAPSD risk genes, which were connected to at least one SCZ risk gene. Given the direct neighbors of MAPSD genes, we categorized them into four groups (**Figure 6d**) followed by counting the number of SCZ risk genes which are connected to each MAPSD risk gene within each group. 93% of MAPSD genes have 1 to 30 direct neighbors among which the median percentage of SCZ risk genes is ∼30%. In other words, on average, 30% of the accumulated signals in MAPSD risk genes were transmitted directly from neighboring SCZ risk genes, while the remaining signal intensities are transmitted from distant genes. This is remarkable given that MAPSD can capture the signals from distant risk loci so that the convergence of small effect size loci can be observed and modeled. Another major property of MAPSD is its resilience against noise. Markov operators in graph-signal processing act as a low-pass filter^68^. Therefore in case of introducing false signals, i.e., noise, to the MAPSD initial signal matrix, these signals will automatically filter out during the signal diffusion. As a result, MAPSD is significantly noise-resistant. MAPSD was able to recover a significant fraction of known SCZ susceptibility genes from multi-omics studies. For example, in a recent study by Wang *et al*^5^., multiple SNPs were reported to be associated with the disease. A significant overlap between MAPSD identified genes and their reported loci was observed (FET P-value=2.1×10^−4^, Enrichment Ratio=3.2). Among them, 85% of these genes were enriched in neuronal cells in cerebral cortex, 7.5% in Purkinje cells in cerebellum, and 7.5% in neuronal cells in caudate. This observation further supports the mechanism introduced in MAPSD to jointly model mutual interactions between omics data modalities for identification of novel risk genes and susceptibility risk modules in PPI networks.

MAPSD takes advantage of high-dimensional omics data and is not tied to specific phenotypes. Therefore, it can effectively be applied to any complex disease such as autism spectrum disorders (ASD) or autoimmune diseases, when necessary multi-omics data sets are available. MAPSD provides an ideal platform to leverage the outcomes of ongoing massive-scale projects such as The Psychiatric Genomics Consortium (PGC)^69^, the largest consortium in psychiatry genetics, and the PsychENCODE project^70^, which is actively generating extensive epigenomic data on various psychiatric disorders. We envision MAPSD to be useful to the community to catalyze integrated evaluation of candidate genes for various neuropsychiatric and neurodevelopmental disorders at a systems level.

## Experimental Procedures

### Description of the data used in the study

Interaction networks used in this study were collected from three sources including: PICKLE 2.3^32,33^, The Human Reference Interactome^31^, and human Interactome Database^30^. Upon removing the duplicate interaction, the final network being used by MAPSD contained 232,801 interactions. The list of differentially expressed genes were obtained from the CommonMind Consortium^2^. GWAS hits on SCZ were downloaded from the CLOZUK consortium^4^ and Psychiatric Genomics Consortium^3^. Rare and *de novo* mutations were downloaded from denovo-db v.1.6.1^25^. DNA methylation data were downloaded from the works by Vitale *et al.*^26^, Aberg *et al.*^27^, and Alelu-Paz *et al*.^28^. Open chromatin accessibility peaks were downloaded from the study by Bryois *et al*.^29^. Protein abundances in all of the tissues and cell-types as well as the subcellular localization of all of the proteins were obtained from the human Protein Atlas project^7,8^. Tissue-specific gene expression levels were obtained from the GTEx project^54^ consortium on 53 tissues.

### Creating the Signal Matrix

The initial signal matrix *S*, is an overlaid column vector which contains the cumulative levels of biological evidences such as transcriptional signatures, methylation, GWAS etc. For each level of information for a specific gene, we add a point 1 if there was an evidence. To create *S*, first we introduce evidence matrix *E*_*G*×*L*_ where *G* denotes the total number of genes and *L* is the number of omics data layers (in this study, 5). Therefore:

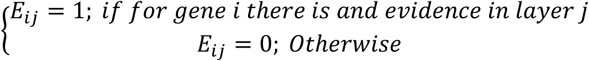

Next, using *E*, we can create *S* as follows: 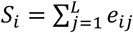 As example if a gene *i* is DE and differentially methylated, then *S*_*i*_ = 2.

### Adjusting the PPI network weights and creating the affinity matrix

Subcellular localization data used in MAPSD were downloaded from Human Protein Atlas project^7,8^. In total, 32 subcellular domains were available. To project this information onto the PPI network, first the affinity matrix *A* was created. *A* is a *n* × *n* binary matrix where *a*_*ij*_ = 1 if two proteins *i* and *j* are connected in the network, otherwise *a*_*ij*_ = 0. *n* denotes the total number of unique proteins in the PPI network. MAPSD scans the entire elements of *A* and checks its localization microdomain. If two proteins *i* and *j* are connected in the network while co-localizing in the same microdomain, then *a*_*ij*_ = 1.5. However, If two proteins *i* and *j* are connected in the network while not being co-localized in the same microdomain, then *a*_*ij*_ = 1. Note that *A* is a symmetric matrix, i.e., *a*_*ij*_ = *a*_*ji*_.

### Creating the Markov Transition matrix from affinity matrix

Upon adjusting the raw affinity matrix to contain the subcellular localization information, MAPSD obtains the Markov operator matrix (*M*). *M* is a *G* × *G* transition probability matrix whose element *m*_*ij*_ denoted the probability of single-step random walk from the node *i* to the node *j* and *G* denotes the total number of proteins being considered. Leveraging Random Walk Laplacian in the Graph Theory^71^, *M* can be obtained as follows: *M* = *D*^−1^*A* where *A* denotes the adjusted affinity matrix above which consists subcellular localization information on all of the edges in the network and *D* represents the degree matrix. *D* is a diagonal matrix of the degree *n*, generated from *A* whose non-zero elements can be obtained as follows: 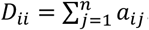. Therefore, each element of the main diagonal in *D* equals the row-wise summation of its corresponding protein in the affinity matrix *A*.

### Creating tissue/cell-specific signal matrix

To use the knowledge on the expression levels of each protein in each cell within each tissue, Human Protein Atlas data was leveraged. In this data, expression levels are defined by four qualitative terms including High, Medium, Low, and Not Detected. To employ his knowledge in MAPSD, we converted them into a weight matrix *W*_*G*×*T*_ where *G* is the total number of proteins being considered and *T* is the total number of tissues and cell-types. The total combinations of tissues and cell-types in this study is 131. Therefore, the expression degree of protein *i* in the tissue/cell *j* is denoted by *w*_*ij*_ as follows:

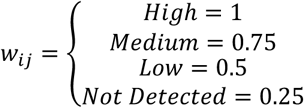

Later, we convert the signal vector *S* to tissue/cell-specific signal matrix *S*^*^ by scalar multiplying the weight matrix *W* and the initial signal vector *S* as follows:

*S*^*^ = *W* ⊙ *S* where *S*^*^ is a *G* × *T* matrix and ⊙ denotes dot (scalar) product. Here, 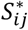 represents the disease signal intensity of the protein *i* in the tissue/cell *j*.

### Signal diffusion process in MAPSD

MAPSD uses the Markov operator matrix *M* and tissue/cell-specific signal intensity matrix *S*^*^ to initiate the diffusion process. During the diffusion process, given the topology of the PPI network, for each combination of tissues and cell-types, signal intensities of SCZ risk loci are propagated onto the network so the signal intensities of unknown proteins are estimated. The higher the signal intensity of a protein in the brain, the higher the likelihood of its association to SCZ. MAPSD is an iterative process where in each iteration signal intensities from disease risk genes are propagated through the network using the following equation: *S*^*t*^ = *M*^*t*^ × *S*^*^ where *t* denotes the diffusion time, i.e., the length of a random walk of size *t* from each node. A critical point to address during the diffusion process is choose of an appropriate diffusion time given that very large values of *t* leads to over-smoothness of the signal intensities. In other words, when the signal are over-smooth, then the signal intensities across all of the network will converge to a constant value leading to the loss of useful information. To avoid this situation, we have created a termination criterion called Smoothness Rate (*R*) as follows: 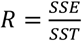 where *SSE* stands for Sum of Square Error and *SST* stands for Sum of Square Total and can be calculated as follows:

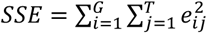 where *e* denotes a single element of the error matrix *E* = *M*^*t*+1^*S*^*^ − *M*^*t*^*S*^*^.

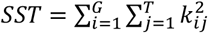 where *k* denotes a single element of the total matrix *K* = *M*^*t*+1^*S*^*^ + *M*^*t*^*S*^*^. MAPSD terminates the diffusion process if *R* ≤ 0.05. In other words, if the normalized difference of changes between signal intensities do not change at a certain threshold, then MAPSD stop the diffusion to avoid over-smoothing the signals of the protein across the network.

### Pathway enrichment analysis

Pathway enrichment and GO analysis were conducted using WebGestalt^72^ v. 2019. KEGG was used as the functional database the list of expressed genes were used as the background. The maximum and minimum number of genes for each category were set to 2000 and 5, respectively based on the default setting. Bonferroni-Hochberg (BH) multiple test adjustment was applied to the enrichment output. FDR significance threshold was set to 0.05.

### Code and data availability

MAPSD scripts and all of the data required for running the platform are available online at: https://github.com/adoostparast/MAPSD.

## Acknowledgements

This study was supported by NIH grant MH108728 (K.W.), Alavi-Dabiri Postdoctoral Fellowship Award (A.D.T.) and CHOP Research Institute (K.W.).

## Author Contributions

A.D.T. conceived the method, coded the algorithm, analyzed the results, and wrote the manuscript. J.D. provided guidance on the study design and the interpretation of results, and revised the manuscript. K.W. conceived the method, supervised the study, and edited the manuscript.

